# Beyond Conventional Models: Recreating the Initiation, Evolution, and Genome of GBM

**DOI:** 10.1101/837138

**Authors:** A Bohm, J DePetro, C Binding, A Gerber, N Chahley, M Ware, K Thomas, S Bukhari, C Chen, E Chahley, C Grisdale, D Berger, S Lawn, Y Yu, R Wong, Y Shen, H Omairi, R Mirzaei, L Maxwell, H Pederson, V Yong, S Weiss, J Chan, P.J. Cimino, J. Kelly, S.J.M. Jones, E Holland, M.D. Blough, J.G. Cairncross

## Abstract

**Background:** Imagining ways to prevent or treat glioblastoma (GBM) have been hindered by a lack of understanding of its pathogenesis. Although PDGF-AA overexpression may be an early event, critical details of the core biology are lacking. Existing PDGF-driven models replicate its microscopic appearance but not the genomic architecture characteristic of the human disease. Here we report a new model of GBM that overcomes this barrier to authenticity.

**Methods:** Using a method developed to study neural stem cells, we investigated the effects of PDGF-AA on subventricular zone (SVZ) cells, the putative cell of origin of GBM. We micro-dissected SVZ tissue from p53-null and wild-type adult mice, established primary cultures in media supplemented with PDGF-AA, and assessed cell viability, proliferation, genome stability, and tumour forming potential.

**Results:** Counterintuitive to its canonical role as a growth factor, we observed immediate and massive death of SVZ cells in PDGF-AA. Wild-type cells did not survive in PDGF-AA. However, a small fraction of null cells evaded apoptosis, displayed attenuated proliferation, gradually accumulated whole chromosome gains and losses, and, signalled by sudden rapid proliferation and growth factor independence, became tumorigenic in immune-competent syngeneic mice. Transformed cells had an OPC-like profile, were resistant to PDGFR-α inhibition, and harboured highly abnormal karyotypes similar to those seen in human GBMs.

**Conclusion:** This model associates genome instability in SVZ cells with chronic exposure to PDGF-AA; it is the first model to replicate the genomic landscape of GBM and first in which the earliest phases of GBM can be directly observed.

**IMPORTANCE OF STUDY:** We have developed a mouse model in which the initiation, evolution and genomic landscape of GBM can be thoroughly studied thus paving the way for ideas about how this deadly brain cancer might be prevented, interrupted at an occult stage, or treated with very different therapies.

## INTRODUCTION

The genomic architecture of the isocitrate dehydrogenase (IDH) wild-type form of GBM, common in older adults, consists of signature chromosomal gains and losses (1). Despite an increasing detailed annotation of these and other molecular alterations, and knowledge of the cellular processes affected by these abnormalities, the prognosis for patients has changed very little, and no new therapies have emerged. In an effort to better understand the origins of IDH wild-type GBM, Ozawa *et al.* used mathematical modeling of TCGA data to identify gain of chromosome 7 with associated overexpression of PDGF-AA as a likely initiating event, often accompanied by losses of CDKN2A, p53, and chromosome 10 (2). Others have also pointed to amplification of PDGF-AA as an early event in the pathogenesis of IDH wild-type GBM (3).

To investigate the interaction of PDGF-AA with other GBM-associated alterations, Ozawa *et al* overexpressed PDGF-AA in p53-null adult mice and generated tumors that were histologically indistinguishable from human GBM. More recently Koga *et al* engineered pluripotent human stem cells to generate human-like GBMs in immune-compromised mice by combining PDGF-AA receptor (PDGFR-α) activation with other alterations in human GBM, including loss of p53, PTEN, and NF1 (4). Further, Jun *et al* produced murine GBMs by constructing mutant mice engineered to overexpress human *Pdgfr-*α in a Cre recombinase dependent manner when crossed with mice carrying a conditional loss of p53 function (5). These models support the view that activation of the PDGFR-α pathway plays a key role in GBM pathogenesis. Each model yields a tumour that is histologically similar to human GBM and captures some of the heterogeneity and clinical drug responses of the human disease, but, to date, none has captured the chromosomal instability and evolutionary trajectory of GBM, which may be important to a deep understanding of disease initiation and to early treatment strategies or risk reduction.

Here, we report a model of GBM in which p53-null and heterozygous SVZ cells transform *in vitro* in PDGF-AA, but not in other factors implicated in the pathogenesis of GBM. Not only do SVZ cells transform, but they also acquire genomic features that are remarkably similar to those of the human disease. Knowing that GBMs likely arise from progenitor cells (6), many of which reside in the subventricular zone (SVZ) of the brain, we prepared SVZ cultures from adult mice using a technique developed to study neural stem cells (7). To simulate oncogenic signalling, cultures were supplemented with epidermal growth factor (EGF) and fibroblast growth factor (FGF), or with PDGF-AA, ligands for aberrantly activated GBM pathways (8). Because the p53 pathway is frequently compromised in GBM (8), we established primary cultures of null and heterozygous cells in different growth factors and monitored their phenotypes compared to wild type cells. We found that cells with p53 compromise behaved differently than wild type cells, and both behaved differently in PDGF-AA than EGF/FGF: only null and heterozygous cells transformed, and only in PDGF-AA. Transformed cells displayed early genomic instability and subsequently generated GBMs in immune-competent syngeneic mice. Using this approach we were able to observe the earliest stages of transformation in cells that evolved to GBMs.

## MATERIALS AND METHODS

### Mice

Eight week-old (p53−/− B6.129S2-Trp53^tm1Tyj^/J and p53+/+ C57BL/6J) mice were purchased from The Jackson Laboratory. All experiments involving mice were conducted in accordance with animal care procedures at the University of Calgary, Protocol #M08029.

### SVZ Neurosphere culture

Neurosphere (sphere) cultures were established in serum-free media as described (9) (NeuroCult™ Media, Stem Cell Technologies, #05700) supplemented with human PDGF-AA (20 ng/mL; Peprotech 100-13A) *or* EGF (20 ng/mL; Peprotech AF-100-15) plus FGF-2 (FGF; 20 ng/mL; Peprotech AF-100-18B) and heparin sulfate (2μg/mL; Stem Cell Technologies 07980). Spheres were passaged at a diameter of 100μm or every 1 to 3 weeks.

### Cell death assay

Cell death was documented using an AnnexinV staining kit (V13241, Life Technologies) as per manufacturer’s instructions. Cells were analyzed on a BD Biosciences LSRII flow cytometer using FACSDiva.

### Proliferation assay

Proliferation was assessed using Click-iT EdU Proliferation Assay (C10634, Life Technologies, USA). 5 × 10^5^ cells were incubated for 2.5 hours in 5-ethynyl-2’-deoxyuridine (EdU) and assessed as instructed by the manufacturer on a BD Biosciences LSRII flow cytometer using FACSDiva (software).

### Syngeneic intracranial implantation

Cells were implanted as described (10). After implantation mice were monitored closely until they began to show signs of illness (weight loss > 15%), at which point they were euthanized. Brains were fixed via cardiac cannulation and perfusion with 4% PFA, or through submersion in 4% PFA overnight.

### Staining and immunohistochemistry (IHC)

IHC was performed as described (11). The antibodies used were: anti-Nestin (1:200; Millipore MAB353), anti-GFAP (1:500; Millipore MAB360) and anti-Olig2 (1:400; Millipore MABN50) with goat anti-mouse IgG-HRP secondary antibody (1:2000; Santa Cruz sc-2005). Peroxidase signal was detected using the ABC Elite kit (Vector Laboratories) and DAB substrate kit (Sigma D4168) as instructed by the manufacturer.

### Phospho-RTK arrays

Mouse Phospho-Receptor Tyrosine Kinase Array Kits (ARY014, R&D Systems) were used to assess receptor phosphorylation, as per manufacturer’s protocol. Arrays were visualized with ECL reagent and Hyperfilm (GE Healthcare).

### Viability assays

Cell viability was assessed in triplicate with the AlamarBlue Viability Assay (Medicorp, USA). Cells were seeded in 96-well plates (5000 cells per well). AlamarBlue reagent was added following treatment and absorbance was measured following 6 hours incubation.

### PDGFR-α inhibition assays

Spheres were dissociated and seeded at 2,000 cells/100μL in a 96-well plate in triplicate. Each well was supplemented with PDGFR-α blocking antibody (1:100; R&D Systems), or Imatinib (Seleckchem). Cultures were incubated at 37°C for six days and viability assessed.

### PDGFR-α knockdown

RNAi specific for murine *Pdgfr-*α was obtained from Invitrogen (Cat# 4390771, ID#s71418). Control scrambled RNAi (Invitrogen, Cat#4390843), *Egfr* RNAi (Catalog #4390771, Lot #ASO216GD, ID: s65372) and transfection reagent Lipofectamine RNAi Max (Qiagen) were used. To transiently inhibit *Pdgfr-*α expression, a modified version of the Manufacturer’s protocol for a reverse transfection was used. Transfections occurred immediately after plating and expression of *Pdgfra* RNA. PDGFRα protein was analyzed 3-7 days later.

### Western blot

Protein was extracted as described (10). Antibodies used were Anti-PDGFR-α (1:1000; Cell Signaling 3174); anti-EGFR (1:2000; Cell Signaling 4267); anti-Olig2 (1:2000; Millipore MABN50); anti-Nestin (1:2000; Millipore MAB353); anti-NG2 (1:1000; Cell Signaling 4235); anti-GFAP (1:1000; Millipore MAB360); anti-β3-Tubulin (1:1000; Cell Signaling 4466); and anti-β-actin (1:5000; Cell Signaling 3700). Secondary antibodies were goat anti-mouse and anti-rabbit IgG-HRP (1:2000; Santa Cruz sc-2005 and sc-2004, respectively).

### Surface marker assessment

Samples were stained for surface markers and incubated in relevant antibodies in the dark (1 hour at 4°C) and analyzed on the BD LSR II flow cytometer. IgG/REA controls were used for gating. Antibodies used: anti-Prominin-1 (CD133)-PE (Milentyi Biotec, 130-102-834); Rat IgG1-PE (Milentyi Biotec, 130-103-042); anti-CD34-FITC (Milentyi Biotec, 130-105-890); REA Control-FITC (Milentyi Biotec, 130-104-626); anti-CD44-FITC (Milenty Biotec, 130-102-933); Rat IgG2b-FITC (Milentyi Biotec, 130-103-088); anti-SSEA-1 (CD15)-PE (Milentyi Biotec, 130-104-989); and REA Control-PE (Milentyi Biotec, 130-104-612).

### Transferring EGF/FGF cultures to PDGF-AA

SVZ cells maintained in EGF/FGF were dissociated and re-plated in media supplemented with PDGF-AA. At each subsequent passage, total and live cell counts and percent viability in Trypan Blue were assessed.

### γH2AX immunoflourescent staining

Staining was done as described (12). Antibodies were anti-gamma H2AX (1:800; Abcam ab26350) and anti-phospho H3 (1:500; Abcam ab47297). Secondary antibodies were anti-rabbit IgG-Alexa Fluor 594 (1:800; Life Technologies A11037) and anti-mouse IgG-Alexa Flour 488 (1:800; Life Technologies A11001). Nuclei were stained with DAPI (1:1000; 1mg/mL stock diluted to 1μg/mL in 1X PBS; Sigma Aldrich D9542).

### Cell cycle

Cells are harvested at indicated times and treated with Accumax to obtain a single cell suspension. Propidium iodide staining was completed as described (13). Cells were analyzed on a FACscan Flow Cytometer (Becton Dickensen) and cell cycle distribution determined using ModFit LT 2.0 software (Verity Software House).

### G-band karyotyping

Karyomax Colcemid^®^ (Cat. #15212-012, Gibco, CA, USA) was added to cells at 80-90% confluence and kept in a CO_2_ incubator at 37°C (1.5 hours). Cells were collected, treated with 100µL Accutase (Cat. #7920, STEMCELL Technologies, BC, Canada) at room temperature (5 minutes), suspended in 8mL 0.075M KCl, incubated in 37°C (15 minutes), and Carnoy’s Fixative added. Cells were then centrifuged at 1000rpm for 10 minutes at room temperature and pellets collected. After three rounds of fixation, cells were re-suspended in fixative and dispersed on glass slides and baked at 90°C (1.5 hours). Routine G-banding analysis was carried out. Twenty metaphases per line were examined.

### aCGH

DNA was isolated using the DNEasy extraction kit (Qiagen) and processed on the Agilent Mouse 1×1M array. Data preprocessing was performed using snapCGH [background correction (method=minimum) and within-array normalization (method=median)]. Within-array replicates were combined into a single value by determining mean value, and data segmented using the DNAcopy implementation of the circular binary segmentation (CBS) algorithm with a default-parameter smoothing and outlier correction (smooth.CNA function). The Integrative Genomics Viewer was used to identify and visualize regions of aberrant copy number.

### Statistical Analysis

Graphpad Prism (Version 8) was used for all analyses. One-way ANOVA was used to assess statistical significance for multiple comparisons within a sample set, while the unpaired t-test was used to assess significance in sample sets of two groups.

## RESULTS

### P53-null SVZ cells proliferate in EGF/FGF but only transform in PDGF-AA

SVZ cultures were prepared as described by Reynolds and Weiss (7). In EGF/FGF, primary SVZ cultures of p53-null cells formed large symmetrical spheres (i.e. diameter > 100μm) within one week (Fig. 1A). Because of their rapid proliferation cultures required weekly passaging (denoted as ‘P’, see methods). The continuous rapid expansion of these cultures led us to inquire whether they had acquired the capacity to proliferate in the absence of EGF/FGF. To test this possibility, null cells from early (P≤4) and later (P≥8) passage cultures were transferred to media without EGF/FGF and immediately stopped dividing (Fig. 1A). Even after 12 months of rapid proliferation in EGF/FGF, null cells remained growth factor dependent. P53 wild type cells behaved identically (Fig. S1A). These findings show that EGF/FGF supports the proliferation of SVZ cells, but continuous signalling does not lead to growth factor independence, even in the absence of p53.

**Fig. 1:**
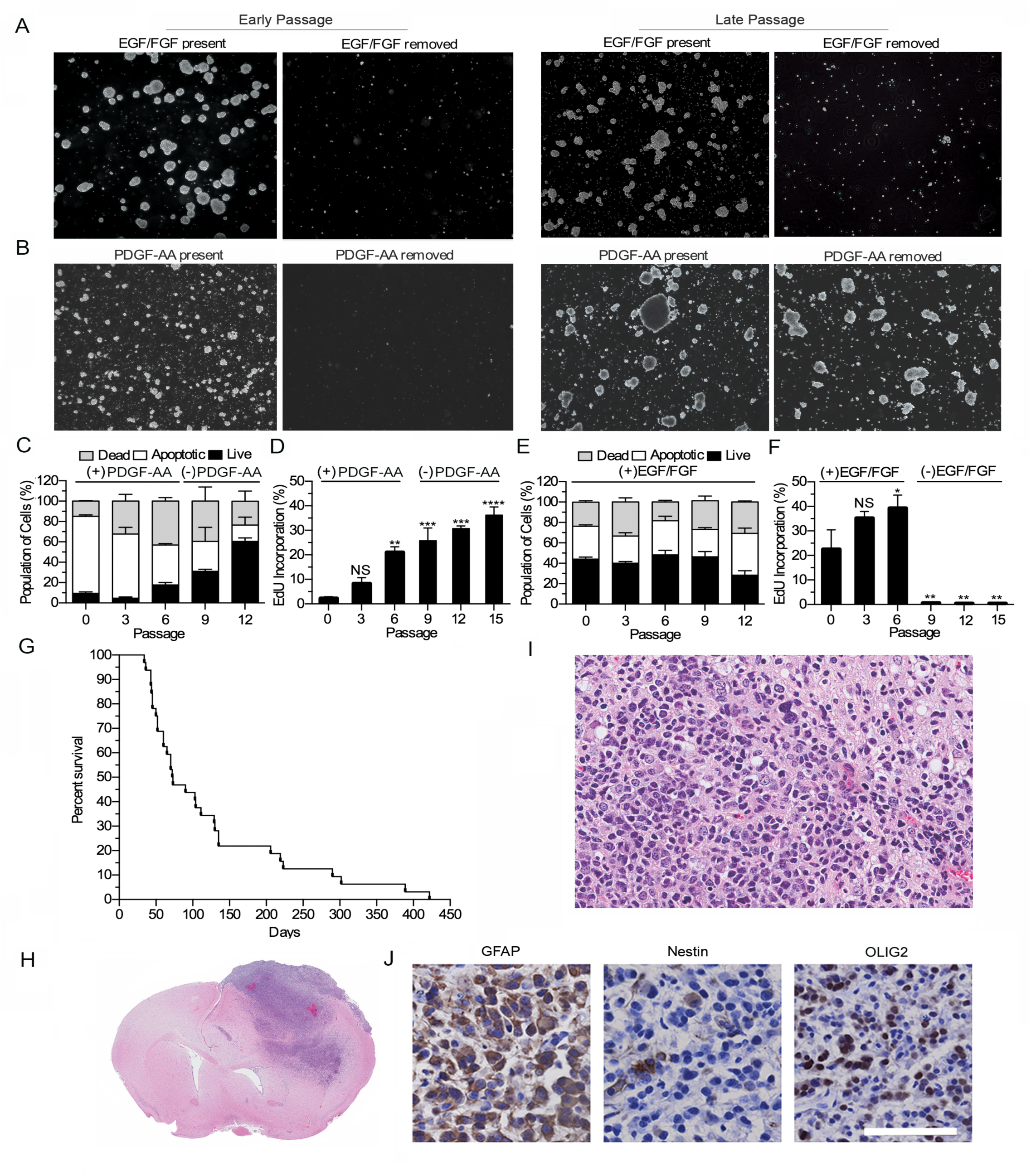
P53-null SVZ cell cultures acquire growth factor independence and become tumorigenic when cultured in PDGF-AA. **(A)** P53-null cells cultured in EGF/FGF form large spheres at early passages (P≤4) but cease proliferating when EGF/FGF is removed (*top left*). At late passages (P≥8), null cells proliferate rapidly and remain EGF/FGF dependent *(top right)*. **(B)** Early passage P53 null cells display attenuated proliferation and sphere formation in PDGF-AA and remain PDGF-AA dependent. At late passages (P≥8), cells rapidly form spheres in presence or absence of PDGF-AA. **(C)** Annexin V staining reveals a high percentage of p53-null SVZ cells (>70%) in early passage PDGF-AA cultures are dead or apoptotic, and that this proportion decreases over time as the live cell fraction increases. **(D)** EdU-labelling of p53-null cultures supplemented with PDGF-AA demonstrates a steady increase in the percentage of SVZ cells undergoing DNA synthesis with increasing passage, and continues to increases further when PDGF-AA independent proliferation is achieved. **(E)** Annexin-V/PI staining reveals that the proportion of dead, apoptotic and live null cells remain constant in EGF/FGF over time. **(F)** EdU-labeling shows that a high percentage EGF/FGF cultured null SVZ cells are replicating their DNA and remain dependent on EGF/FGF supplementation. **(G)** PDGF-AA independent P53-null cells form GBMs in p53 wild type. Tumor bearing mice (n=32) had a median survival of 73 days. **(H & I)** H&E staining reveals typical GBM features including high cellularity, brain infiltration, nuclear pleomorphism, hemorrhage and necrosis. **(J)** Immunohistochemistry reveals presence of GBM markers, including GFAP, Nestin and Olig2 positive cells (scale bar = 50 μm). (NS = p>0.05; * = p<0.05; ** = p<0.01; *** = p<0.001; **** = p<0.0001)

In contrast, primary cultures of null cells in PDGF-AA immediately underwent extensive cell death and displayed markedly attenuated proliferation. Passaging of null cells was not required until three weeks after set-up at which point small irregular spheres and single cells were seen. Early passage PDGF-AA cultures (i.e., P≤4) contained abundant debris and some viable cells, suggesting that proliferation and cell death were occurring simultaneously (Fig. 1B). Around P-8 (i.e., after 100 days) null cells underwent a dramatic change in phenotype and began to divide rapidly, requiring frequent passaging. At this juncture, cells quickly formed large asymmetric spheres, which led us to inquire whether growth factor independence had been achieved. Cells from early (P≤4) and later (P≥8) passages were transferred to media that did not contain PDGF-AA. Early passage cultures ceased to proliferate in the absence of PDGF-AA whereas later passage cultures continued to expand (Fig. 1B). Unlike null cells, wild type cells underwent extensive cell death and did not survive or proliferate in PDGF-AA (Fig. S1B).

To further document the behaviour of null cells in PDGF-AA, we used Annexin V and propidium iodide (PI) staining to measure the fraction of dead, apoptotic, and live cells over time. We observed that <10% of cells were viable at passage 0 (P-0) *versus* 60% at P-12 (Fig. 1C), suggesting the gradual selection of cells able to survive in PDGF-AA. We also used EdU labelling to assess rates of DNA synthesis in PDGF-AA and found that proliferation increased from P-0 to P-15 (Fig. 1D), including after P-8 when cultures displayed PDGF-AA independent growth, again implying the selection of cells capable of proliferating in PDGF-AA. In contrast, the proportions of dead, apoptotic and live cells, and their rates of proliferation as measured by DNA synthesis, remained constant in EGF/FGF (Fig. 1E, F). Markedly attenuated proliferation followed by a period of intense selection culminating in the emergence of a population of rapidly dividing cells, was a phenotype that was only seen in p53-null cells in PDGF-AA. Moreover, this unique PDGF-AA phenotype was not seen in null cells grown in PDGF-AA plus EGF/FGF.

These findings show that PDGF-AA has a counterintuitive anti-proliferative effect on SVZ cells. Rather than rapid growth, we noted cell death, attenuated proliferation, and gradual progression to a PDGF-AA independent proliferative state. These events were seen in multiple independent experiments (n=40) and two of 12 heterozygous p53 cultures in PDGF-AA; heterozygous cells took approximately 10 months to become growth factor independent, and PFGF-AA independent proliferation was associated with loss of p53 protein expression (Fig. S1C, D).

The capacity of p53-null SVZ cells to acquire growth factor independence in PDGF-AA prompted us to inquire whether they had become tumorigenic. Cells from multiple PDGF-AA independent null cell cultures were implanted into the striatum of 8-12 week old immune-competent mice. All mice developed infiltrating high-grade astrocytomas (i.e., GBMs) and were euthanized at the first sign of illness; time to illness ranged from 50 to 420 days (median 73 days), a reflection of their heterogeneous aggressiveness (Fig. 1G & H). Hematoxylin and eosin (H&E) staining revealed cellular neoplasms characterized by frequent mitoses, areas of hemorrhage, and necrosis (Fig. 1I). Tumour cells expressed GFAP, Nestin and OLIG2 (Fig. 1J). Additionally, these high-grade astrocytic gliomas contained small numbers of CD3-positive T-cells and Iba1-positive macrophages and microglia (Fig. S1E, F).

### Transformed cells display phosphorylation of multiple receptor tyrosine kinases (RTKs) and PDGFR-α independent proliferation

The capacity of p53-null SVZ cells to evolve to a growth factor independent and tumorigenic state in PDGF-AA led us to examine whether PDGFR-α was phosphorylated in the absence of exogenous PDGF-AA. Using phospho-RTK arrays (Fig. 2A), we saw two distinct patterns of RTK phosphorylation in 15 independently derived tumorigenic lines (Fig. S2A): i) phosphorylation of PDGFR-α alone (Fig 2A, panel I) and ii) phosphorylation of PDGFR-α together with the EGF receptor (EGFR; panel II), EGFR with AXLR (panel III), or EGFR with the hepatocyte growth factor receptor (HGFR; panel IV). To ensure these results were not an *in vitro* artifact, transformed lines were grown intracerebrally in immune-competent mice and the analysis repeated on tumors; PDGFR-α phosphorylation and similar secondary patterns were seen (Fig. 2A). While further work is needed to clarify which RTKs drive the proliferation of transformed cells and which sites are phosphorylated, these results are intriguing because activation and up-regulation of the EGFR, AXLR, and HGFR have been reported in human GBM and implicated in its pathogenesis (14–16).

**Fig. 2:**
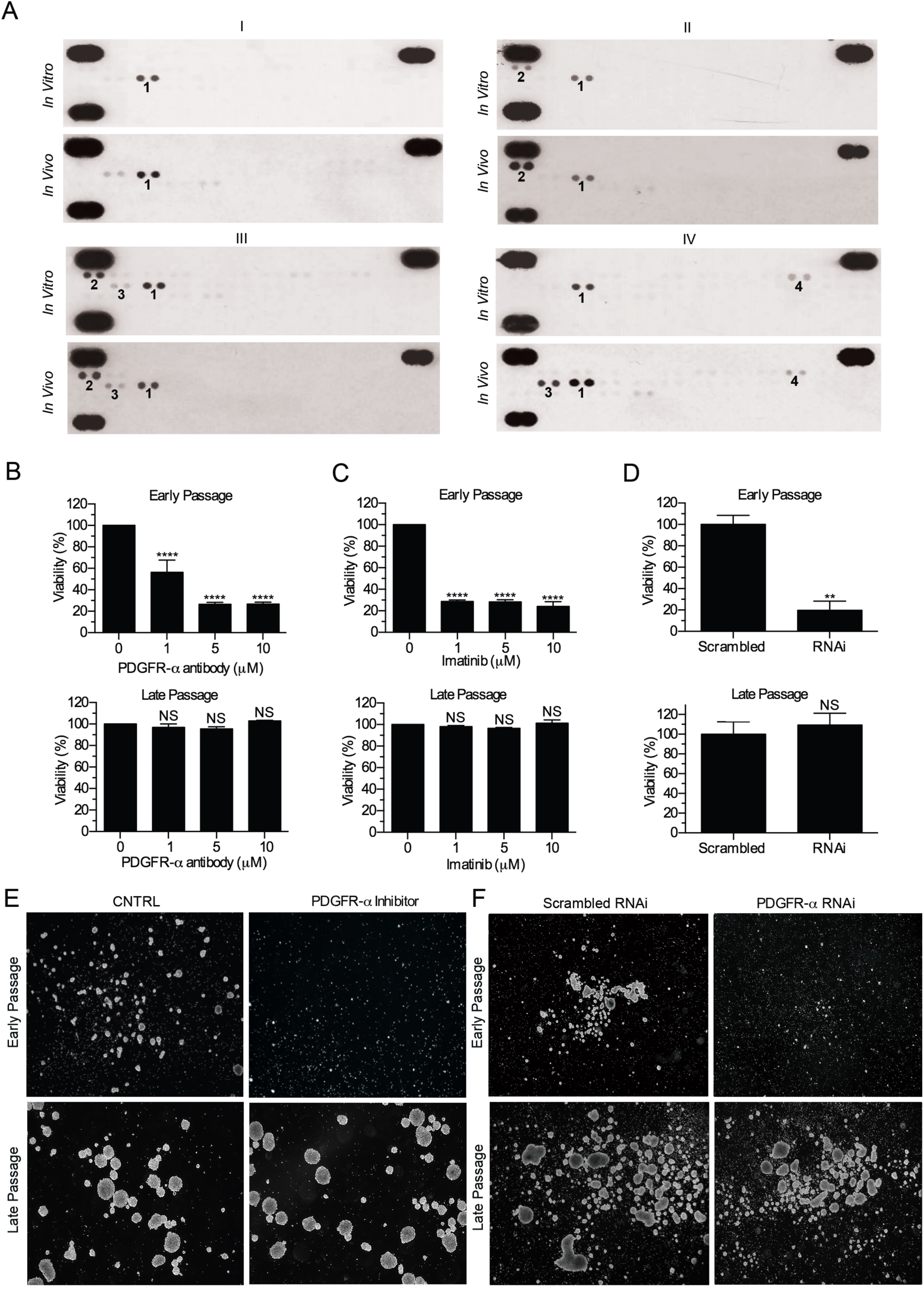
Transformed cells have complex and variable patterns of RTK phosphorylation and are unaffected by PDGFR-α inactivation. **(A)** In PDGF-AA independent transformed cell lines and derived intracerebral xenografts, multiple patterns of RTK phosphorylation were observed including: PDGFR-α alone (*I*); PDGFR-α with EGFR (*II*); PDGFR-α with EGFR and HFR (*III*); *PDGFR-*α with HGFR and AXLR [*1=PDGFR-*α*, 2=EGFR, 3=HFGR, 4=AXLR*] **(B, C, D)** Antibody-mediated inhibition of PDGFR-α (**B**), treatment with a PDGFR-α Imatinib **(*C*)**, and knockdown of the PDGFR-α **(*D*)** decrease cell viability in early passage cultures but do not effect late passage PDGF-AA independent cultures. **(E, F)** Blocking PDGFR-α, or knocking down PDGFR-α leads to a decrease in cell proliferation and sphere formation in early passage p53-null SVZ cultures but not in late passage transformed cultures. (NS = p>0.05; * = p<0.05; ** = p<0.01; *** = p<0.001; **** = p<0.0001)

Persistent PDGFR-α phosphorylation in the absence of PDGF-AA prompted us to ask whether PDGFR-α was required for growth factor independence and necessary to sustain the proliferation of transformed cells. Accordingly, we inhibited PDGFR-α before and after transformation using antibody-mediated blockade, Imatinib, RNAi knockdown, and CRISPR knockout. Receptor blockade, Imatinib, and RNAi decreased SVZ cell viability, proliferation, and sphere formation in early passage PDGF-AA dependent cultures, but had no effect on transformed cells (Fig. 2B-F). Likewise, *Pdgfr-*α CRISPR knockout did not alter proliferation; transformed cells lacking *Pdgfr-*α retained the ability to proliferate in the absence of PDGF-AA (Fig. S2B). Although PDGFR-α was required for progression to a growth factor independent state, these results illustrate that the receptor becomes redundant after transformation.

### Transformed cells have an oligodendrocyte progenitor cell (OPC) profile, proliferate despite DNA damage, and evade apoptosis

The capacity of p53-null SVZ cells to transform in PDGF-AA prompted us to ask whether PDGF-AA selected for a cell that was prone to transform. To assess lineage profiles, we examined Olig2, GFAP, B3-Tubulin, Nestin and NG2 protein expression by Western blotting in PDGF-AA and EGF/FGF cultures (Fig. 3A). Except for the progenitor cell marker, Nestin, which increased over time in PDGF-AA, lineage markers were similar in both conditions. Weak GFAP and β3 Tubulin expression combined with strong NG2 and Olig2 expression was consistent with an OPC profile (17, 18). To further characterize PDGF-AA cultures, the presence of stem cell markers expressed by human GBMs was examined (Fig. 3B). Four transformed lines (I-IV, Fig. 3B) showed differing levels of CD15, CD34, CD44 and CD133 with variable expression; similar heterogeneity is seen in human GBM (19–21).

**Fig. 3:**
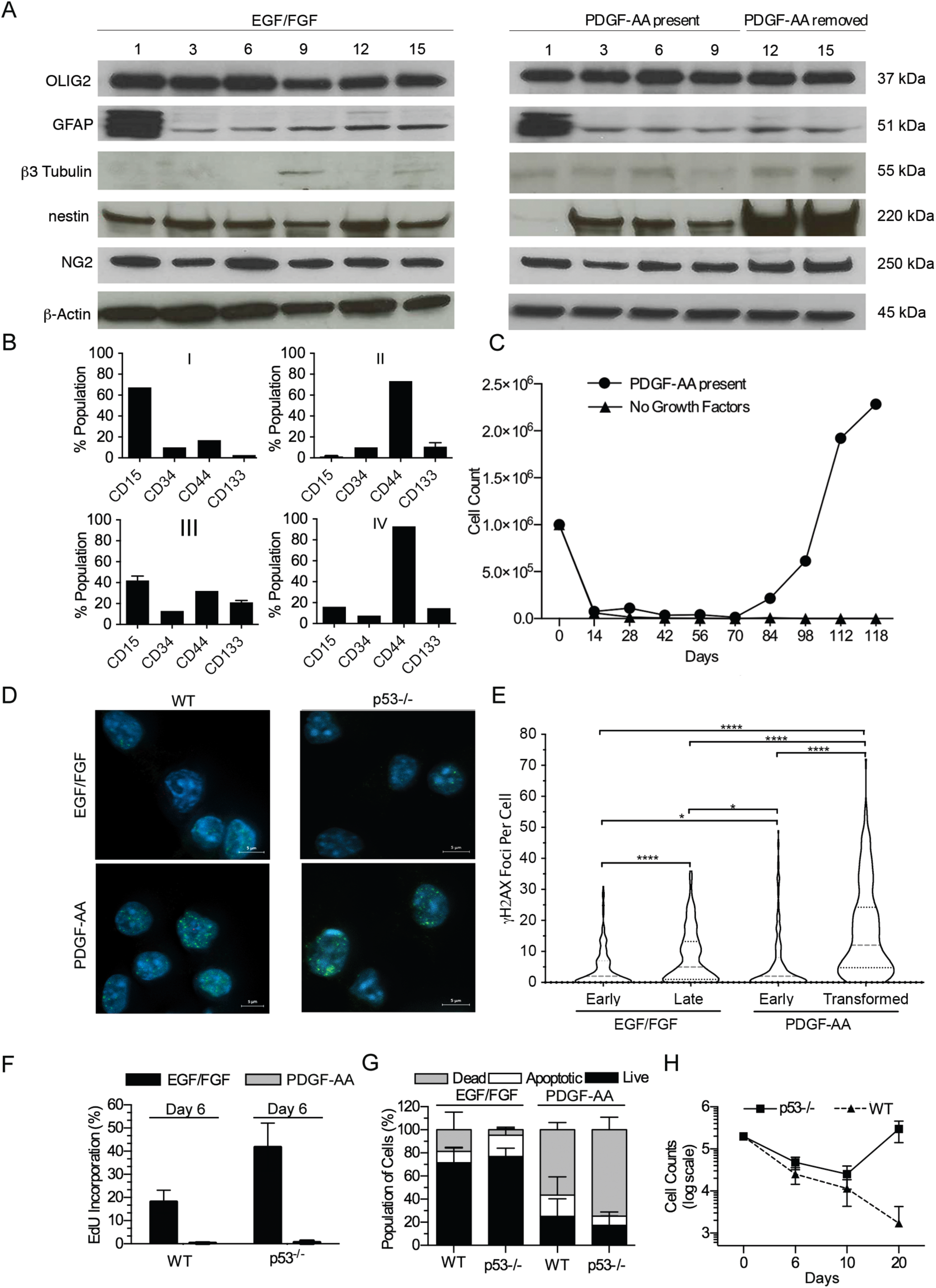
Transformed p53-null SVZ cells have an OPC lineage profile, proliferate in the presence of DNA damage, and evade apoptosis. **(A)** Protein expression of neural lineage markers shows similar profiles in cultures supplemented with EGF/FGF or PDGF-AA. While most markers remain consistent, Nestin expression increases in the PDGF-AA condition. (**B**) Stem and cancer related marker analysis reveal heterogeneous expression of CD15, CD33, CD44 and CD133 in four independent lines (I-IV) transformed in PDGF-AA. **(C)** Null cultures established in EGF/FGF retain the capacity to transform when transferred to PDGF-AA, but are not viable in the absence of growth factors. **(D)** γH2AX staining reveals an increase in the number of DNA damage foci in PDGF-AA cultured cells compared to EGF/FGF, an effect seen in both p53 null and wild type cells. **(E)** Counting γH2AX foci in null SVZ cells cultured in EGF/FGF or PDGF-AA reveals a significant increase in the number of foci per cell, proportional to the time in culture and related directly to PDGF-AA. **(F)** P53 wild type and null cultures stop synthesizing DNA six days after being transferred from EGF/FGF to PDGF-AA. **(G)** P53 wild type and null cells undergo markedly increased apoptosis when transferred from EGF/FGF to PDGF-AA. **(H)** Trypan blue exclusion demonstrates that p53 wild type and null cells undergo decreased proliferation in PDGF-AA but after 10 days the null cells resume proliferating. (NS = p>0.05; * = p<0.05; ** = p<0.01; *** = p<0.001; **** = p<0.0001)

The comparable lineage marker profiles in PDGF-AA and EGF/FGF led us to ask whether cultures established in EGF/FGF retained the capacity to become growth factor independent and tumorigenic in PDGF-AA. To test this possibility, p53-null cells that had been maintained in EGF/FGF for 12 months were switched to PDGF-AA. Control cultures ceased proliferating when EFG/FGF was removed from the media, whereas those transferred to PDGF-AA entered a period of massive cell death and markedly attenuated proliferation (Fig. 3C). As seen in primary cultures of null cells in PDGF-AA, rapid proliferation and sphere formation occurred approximately 100 days later accompanied by PDGF-AA independent proliferation (Fig. 3C) and tumour formation in mice. These findings reveal that p53-null SVZ cells retain the potential to evolve to a growth factor independent state despite prolonged culturing in EGF/FGF, suggesting that exposure to PDGF-AA is the key determinant of progression to growth factor independence. Having observed extensive cell death in SVZ cells in PDGF-AA, and transformation of null cells, effects not seen in EGF/FGF, we inquired whether DNA damage was occurring in the PDGF-AA growth condition, reasoning that damage might trigger both apoptosis and cancer formation (22). To assess DNA strand breakage, we counted γH2AX foci in null and wild type cells in PDGF-AA and EGF/FGF and found greater numbers of foci in PDGF-AA, irrespective of genotype (Fig. 3D & Fig S3A, B). Then, to assess the effect of time in culture on strand break abundance in EGF/FGF versus PDGF-AA, γH2AX foci were counted in early and late passage null cells (Fig. 3E). In EGF/FGF, the mean number of foci in early passage cells was 4.6, which increased to 8.2 in later passage cells (p<0.0001), while in PDGF-AA supplemented cultures the mean number of foci increased from 6.30 to 16.4 (p<0.0001). These studies show that DNA strand breaks increase over time in both growth conditions but are more numerous in PDGF-AA.

The ability of p53-null SVZ cells to proliferate despite DNA damage suggested that null cells in PDGF-AA might be unable to activate cell cycle arrest or apoptotic functions after strand breakage due to loss of p53-dependent DNA damage checkpoint(s). To explore this possibility, cell death and DNA replication were assessed in wild type and null SVZ cells after six days of exposure to PDGF-AA. We observed that extensive cell death had occurred and DNA replication had ceased in cells of both genotypes (Fig. 3F & G). Indeed at day 10, there were equally few viable wild type and null SVZ cells in the PDGF-AA growth condition (Fig. 3H). Thereafter, however, the culture phenotypes diverged. The number of wild type cells continued to decline, whereas the number of null cells increased. After 20 days in PDGF-AA, there were no viable cells in the wild type cultures whereas the null cultures had repopulated. These results demonstrate that absence of p53 allows a small population of null cells with strand breaks to evade apoptosis, proliferate in the presence of PDGF-AA, and transform.

### Chromosome instability occurs in SVZ cells in PDGF-AA

The capacity of p53-null SVZ cells to tolerate DNA damage, become growth factor independent, and tumorigenic led us to search for evidence of genomic instability, because instability is a cancer predisposing mechanism (22). G-band karyotyping was used to assess ploidy, chromosomal integrity, and clonality in multiple SVZ cultures over time. In early passage p53-null cultures in PDGF-AA a small percentage of cells (15%) were observed to be polyploid (Fig. 4A). Clonal chromosomal alterations were infrequent in early passage cultures but losses of chromosomes were seen, especially 12 and 14. Later passage cultures had increasingly complex karyotypes with mosaic and near-tetraploid forms (Fig. 4B). By P-7, 90% of metaphase spreads were tetraploid, and by P-18, 100% were polyploid with clonal and sub-clonal populations (Fig. 4B). Furthermore, by P-18, 90% of cells had lost chromosome 2, 85% had lost chromosome 12, and 100% had lost the Y chromosome; moreover, 60% had gained at least one copy of chromosome 11. In contrast, greater than 80% of SVZ cells in EGF/FGF at all passages had near-diploid karyotypes. Although occasional gains of chromosomes were seen in EGF/FGF, there were no chromosomal losses (Fig. 4C & D).

**Fig. 4:**
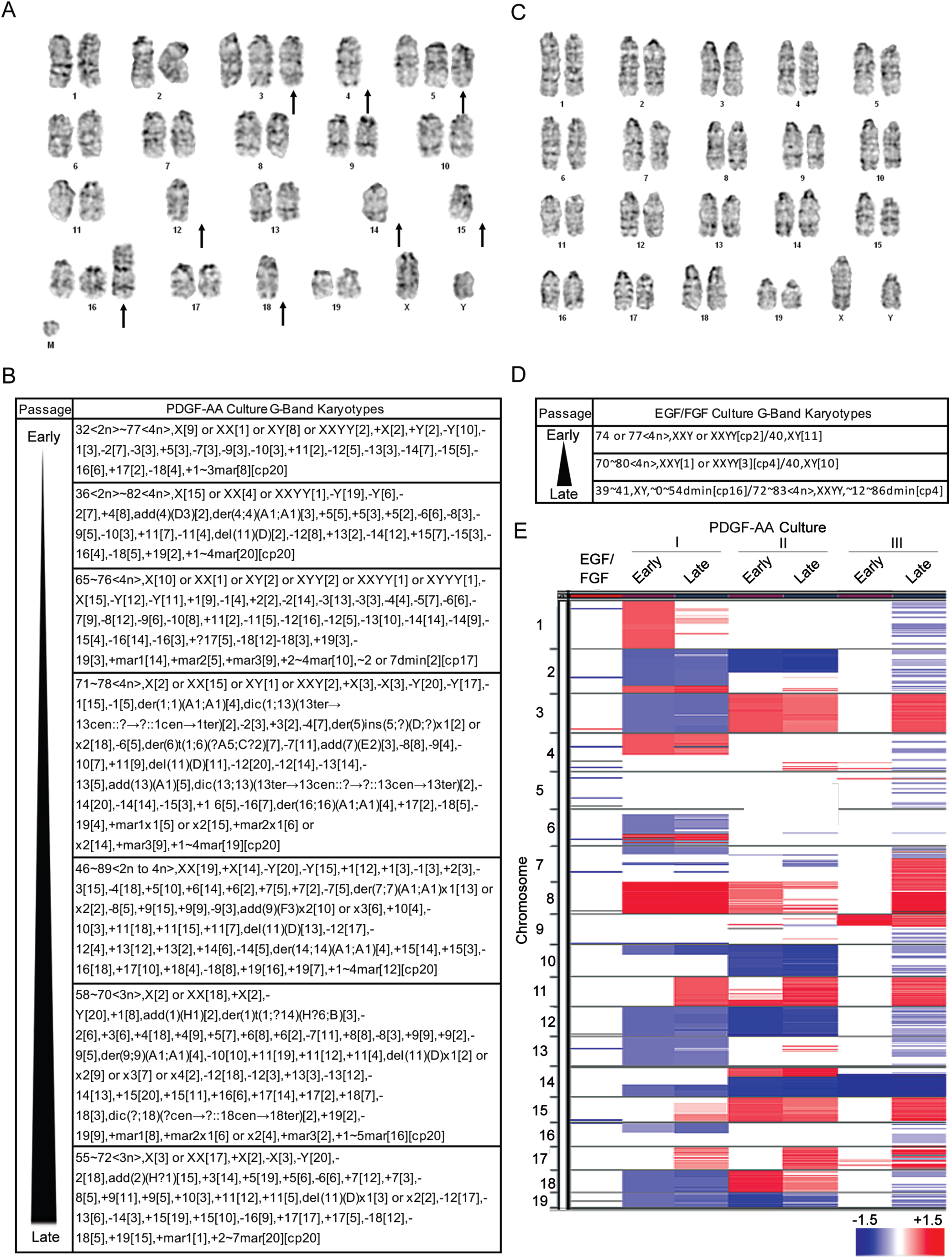
*P53*-null SVZ cells that transform in PDGF-AA have chromosomal instability. **(A)** G-band karyotyping of early passage p53-null cells cultured in PDGF-AA reveal heterozygous loss and gains of chromosomes (arrows). **(B)** G-band karyotypes of early (P4) to late (P18) passage p53-null cultures are shown. Transformed cells display chromosomal alterations that are present early on and become increasing complex. **(C)** G-band karyotyping of p53-null EGF/FGF cultures reveals relatively normal chromosomal number and structure. **(D)** P53-null EGF/FGF cultures have relatively stable karyotypes over time. **(E)** aCGH of an early passage p53-null cells in EGF/FGF versus three pairs of early passage PDGFA-dependent null cells and matched cell lines that have transformed in PDGFA. Cells in EGF/FGF have few alterations whereas null cells in PDGF-AA have many large chromosomal alterations that increase over time in culture.

To better document the nature of these chromosomal alterations, high-resolution array comparative genomic hybridization (aCGH) was performed on a single p53-null EGF/FGF SVZ culture and three matched early and late passage PDGF-AA cultures. As predicted by their karyotypes, p53-null cells maintained in EGF/FGF had no gross chromosomal alterations (Fig. 4E). However, major chromosomal changes including whole and partial gains and losses were seen in the PDGF-AA condition (Fig. 4E). Although some alterations were cell line specific, recurring changes were seen, including losses of portions of chromosomes 2, 10, 12, and 14, and gains of 8, 11, and 15. These regions of recurrent loss are syntenic to multiple areas of the human genome, including chromosome 13, the site of *RB1* in humans, a pathway frequently disrupted in GBM (23), and nearly all of chromosome 10, a pathognomonic deletion in IDH wild type, primary GBM (1, 8). These data show that P53-null cells that transform in PDGF-AA have unstable genomes, a defining feature of human GBM, not seen in other laboratory models.

## DISCUSSION

The earliest stages in the initiation of IDH-wild type GBM are inaccessible to study, hidden within a molecularly heterogeneous disease that often develops explosively with no obvious *in situ* or low-grade stage. Here we address this obstacle to understanding the human disease by developing an *in vitro* system in which the initiation of a GBM can be studied in mice. To our knowledge, this is the first model of GBM that replicates the genomic landscape of the human disease, including recurring chromosomal gains and losses.

Using this model, we uncover a dichotomous role for PDGF-AA in the initiation versus maintenance of GBM. We demonstrate that PDGFR-α is necessary to transform p53-null SVZ cells that are cultured in PDGF-AA, but then discover that the receptor is not required to maintain their oncogenic phenotype. Inactivation or knockout of PDGFR-α does not alter the proliferative capacity of SVZ cells that have transformed in PDGF-AA, nor does it affect their potential to form GBMs in the brains of immune-competent mice. This observation implies that some initiators of malignant transformation will be poor therapeutic targets. Indeed, this finding is consistent with clinical experience: although PDGF-AA overexpression has been implicated in the pathogenesis of many GBMs, these cancers are resistant to drugs that inhibit PDGFR-α signaling (24). We also use this model to show that rapid proliferation of null SVZ cells, a phenomenon often assumed to lead to the accumulation of genetic alterations that transform cells, is insufficient to initiate GBM. P53-null cells divide rapidly in EGF/FGF supplemented media but retain near normal karyotypes and genomes, and do not transform. This counter-intuitive result is consistent with the observation that EGF overexpression and receptor activation are insufficient to generate GBMs in mice (25, 26), suggesting that amplification of EGFR may not be an initiating event in this cancer. Our finding that cells that have transformed in PDGF-AA show secondary RTK activation may be a clue to the origins of EGFR amplication in GBM.

The most remarkable finding emerging from our work is the demonstration that chronic exposure of SVZ cells with p53 compromise (i.e., null or heterozygous) to a ubiquitous growth factor leads to massive apoptosis, attenuated proliferation of selected cells, and their abrupt transformation 100 days later. Although the mechanisms that underlie these responses remain to be fully elucidated, we found evidence of chromosome instability in p53-null SVZ cells that was unique to the PDGF-AA condition. Indeed, one of the earliest alterations in pre-transformed early passage null cells were recurrent losses of specific entire chromosomes or large portions of chromosomes. High levels of DNA damage were found in null cells especially after prolonged exposure to PDGF-AA, but stand breaks were not unique to this condition. These observations introduce the possibility that chromosome segregation errors associated with exposure to PDGF-AA may underlie the malignant transformation of stem-like cells in which p53 or its pathway has been compromised. Indeed, segregation errors can induce mitotic arrest (27), DNA breaks (28) and apoptosis (27) and lead to gains and losses of whole chromosomes (29), features that have been documented in our system. Intriguingly, the areas of recurrent losses in our model harbor GBM-associated tumour suppressor genes.

One final point merits comment. The seemingly abrupt change from PDGF-AA dependent to independent growth, signalled by rapid sphere growth and tumour formation, is reminiscent of the behaviour of IDH-wild type GBM, a cancer that often appears suddenly. The monophasic presentation of this subtype of GBM belies its vast molecular heterogeneity at diagnosis, a paradox suggesting that occult genome instability without visible tumour growth is a feature of the earliest stages of this cancer. In our model, a period of attenuated proliferation in PDGF-AA precedes the rapid expansion and transformation of p53-null SVZ cultures. Perhaps our system can be a tool for understanding and intercepting the prodromal phase of primary GBM.

---

**Fig. S1:**
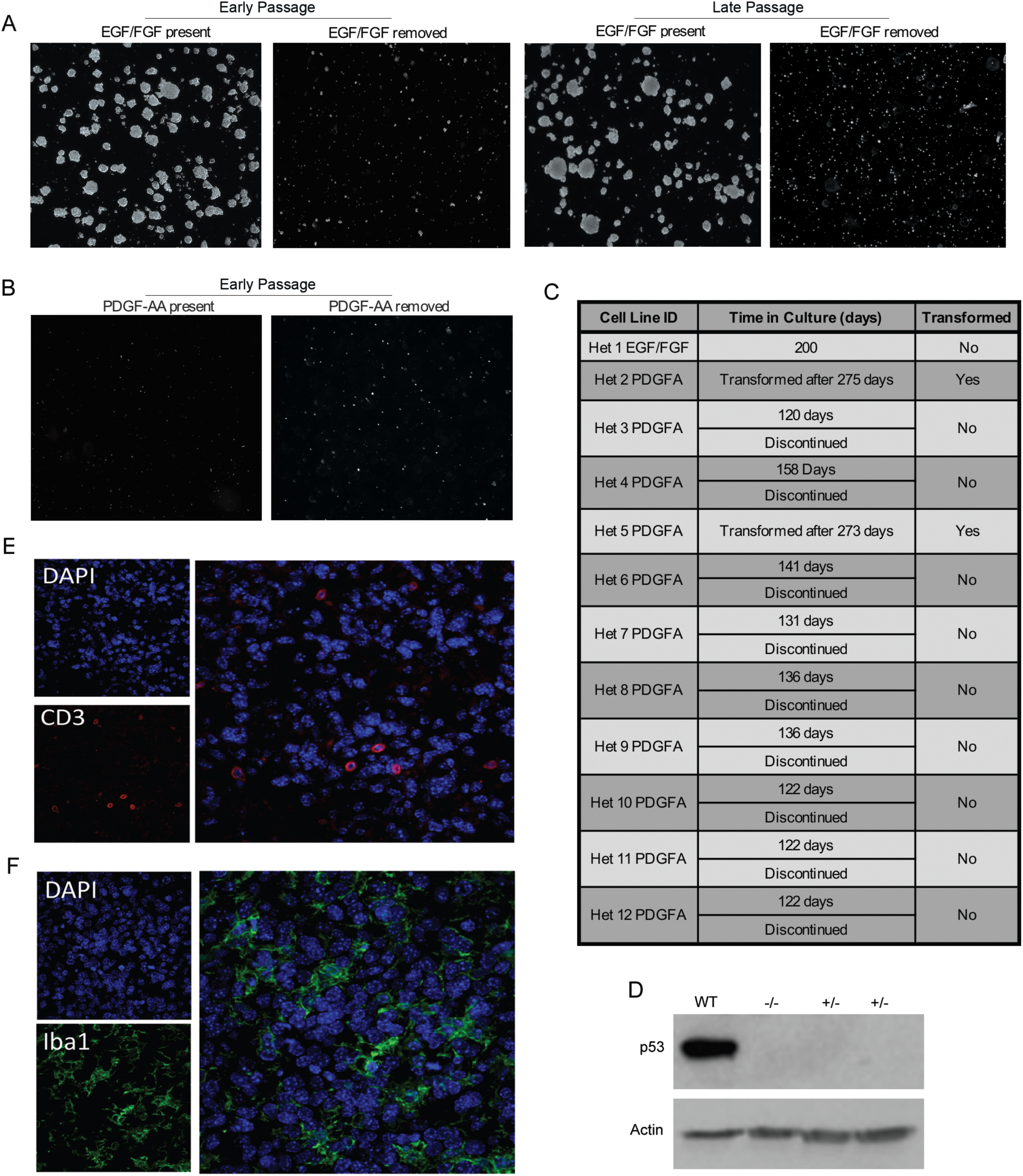
**(A)** P53 wild type SVZ cells proliferate rapidly and form neurospheres in EGF/FGF but remain growth factor dependent, regardless of time in culture. **(B)** P53 wild type SVZ cells do not proliferate in PDGF-AA, regardless of time in culture, and do not become growth factor independent. **(C & D)** Two of twelve cultures heterozygous for p53 (p53-het) eventually became growth factor independent in PDGF-AA. **(E & F)** Staining of tumours from transformed cells for CD3 (T cell) and Iba1 (microglia) show a T cell infiltrate and microglia at the edge of the tumor.

**Fig. S2:**
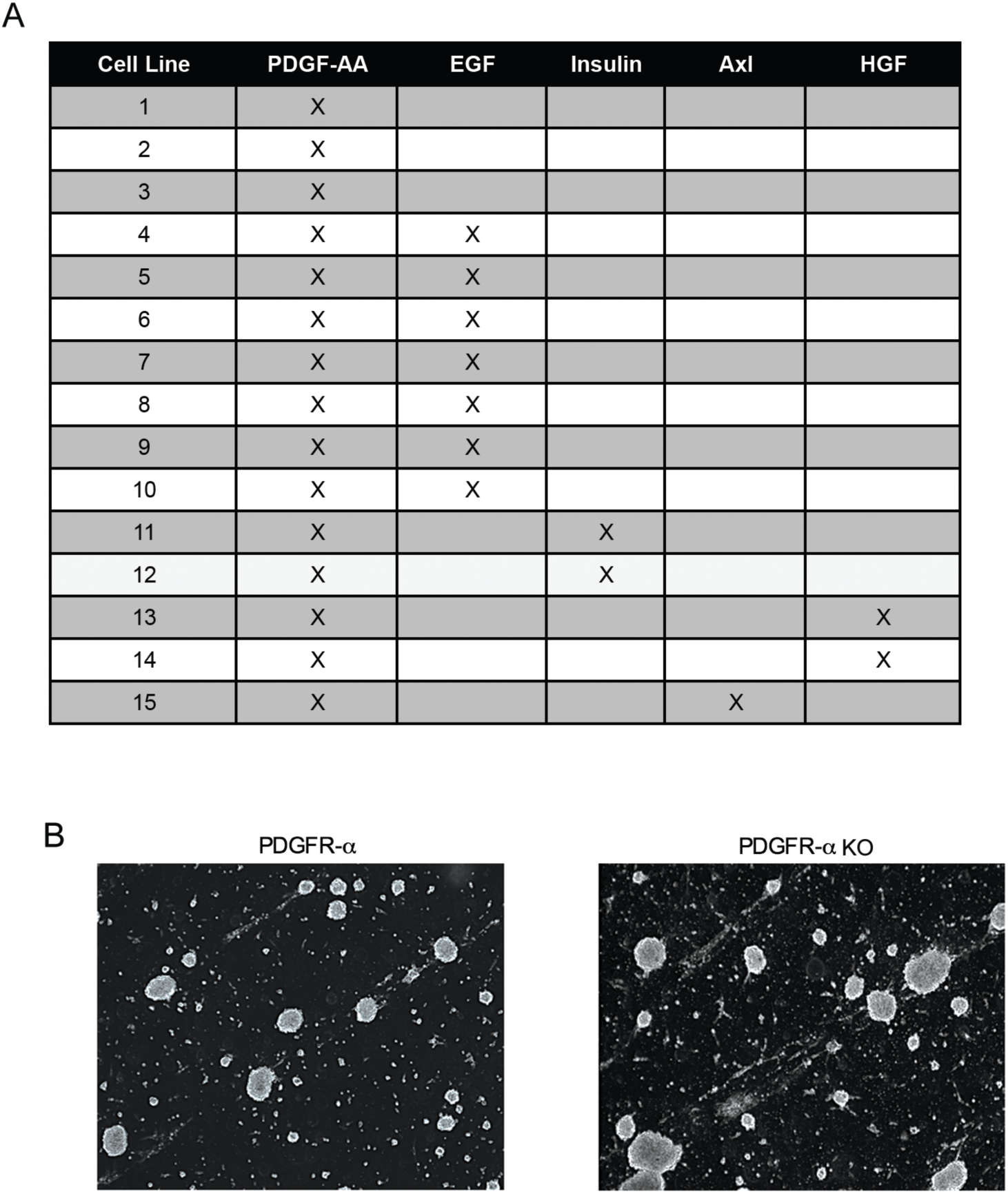
**(A)** Assessment of receptor phosphorylation in 15 PDGF-AA transformed p53-null cell lines reveals PDGF-AA remains phosphorylated in all lines, either alone or in combination with other receptors. [X = phosphorylation detected] **(B)** CRSPR-mediated knockout of PRGFR-α in a PDGF-AA transformed line confirms the receptor becomes redundant after transformation; cell proliferation and neurosphere formation remain unchanged after PRGFR-α knockout.

**Fig. S3:**
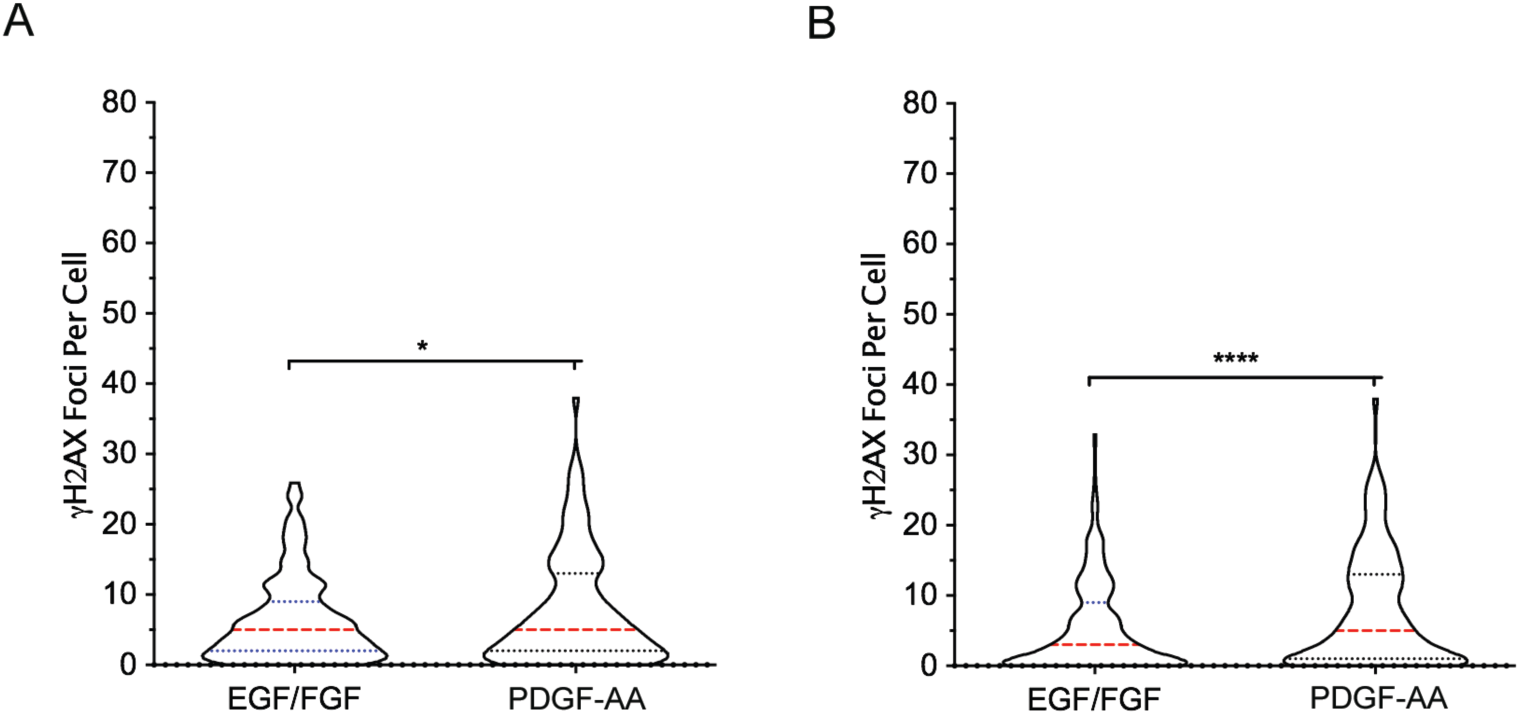
**(A)** Transferring p53 wild type EGF/FGF cultures to PDGF-AA results in a significant increase in γH2AX foci. **(B)** Transferring EGF/FGF p53-null cultures to PDGF-AA also results in a significant increase in γH2AX foci.

## SUPPLEMENTARY METHODS

### Pdgfr-α CRISPR Gene Knockout

pX458 (pSpCas9(BB)-2A-GFP) vector was a gift from Feng Zhang (Addgene plasmid 48138). Short guide sequences which targeting *Pdgr*α exon three are 5’ AAGGAATCGGTCATCCCGAG 3’ and 5’ TAACCTTGCACAATAACGGG 3’, targeting *Pdgfr*α exon six are 5’ ACCCGACGCCCAGGATATCG 3’. All short guide sequences oligonucleotides were synthesized, annealed and cloned into pX458 vector and were confirmed by DNA sequencing at DNA laboratory, University of Calgary. *Pdgfr*α short guide RNA constructs 1, 2, and 3 were transfected with Lipofectamine 2000 (Invitrogen) into a *p53* null SVZ cell line that had transformed in PDGFA as recommended by the manufacturer. Genomic DNA from wild type cells and null cells transfected with Guide 1, 2 and 3 cells was isolated using the KAPA Express Extract Kit (Kapa Biosystems) according to the manufacturer’s instructions. The primers used to amplify genomic region of *Pdgfr*α exon three are: forward 5’ TGGTGGTGAAGCTAACGATG 3’ and reverse 5’ AAACTCGCTGGTCTTGAACG 3’ and *Pdgfr*α exon six are: forward 5’ CTTCGTCGAGATTGAGCCCA 3’ and reverse 5’ CTCGTCAGGCTGCAATGTTA 3’. Surveyor nuclease mutation detection assay was performed using SURVEYOR Mutation Detection Kit (Transgenomic) according to manufacturer’s protocol. After transfection, GFP containing cells were sorting into 96 wells plate by Flow Cytometry. Single cells were then expanded for further analysis (i.e., Western blot and DNA sequencing). All CRISPR gene knockout clones were screened by Western blot using both mouse anti-PDGFRα (Santa Cruz) and rabbit anti-PDGFRα (Cell Signaling) antibodies. The membrane was also probed with Mre11 antibody (Novus) as a loading control. DNA fragments of *Pdgfr*α knock out clone G2-1 and G3-8 around the short guide RNA site of *Pdgfr*α were amplified by PCR, subcloned into pEGFP-C2 vector (Clontech) and plasmid DNA from individual clones were sent for Sanger sequencing at the University of Calgary to confirm indels.

### Iba1 and Cd3 Immunofluorescent Staining

A Leica RM2135 Microtome was used to prepare paraffin-embedded tumor tissue sections. After de-paraffinization and antigen retrieval with Tris-EDTA buffer (10mM Tris Base, 1mM EDTA solution, pH 9), sections were blocked for 1 hour with a solution of 10% horse serum, 1% BSA, 0.1% Triton X-10, and 0.05% Tween 20 in 1XPBS. Then, tissues were subjected to antibody staining with anti-CD3 (Abcam, cat. No.11089, Clone CD3-12, 1:200) and anti-Iba1 antibody (Wako Cat. No. 019-19741, 1:1000) overnight at 4°C. Primary antibodies were diluted in a solution of 1% BSA and 0.5% Triton X-100. After five washes in PBS at room temperature, Alexa 488-conjugated anti-rabbit or Alexa 546-conjugated anti-rat secondary antibodies diluted (1:400; Jackson ImmunoResearch Laboratories) along with nuclear yellow (1:1000) were added for 90 minutes. Sections were mounted in Fluoromount-G (ThermoFisher Scientific) and cover-slipped. Tissues were examined using the Olympus Fluoview FV10i confocal microscope. Images were captured using a 60X objective.

